# miRNA-dependent poly(A) length control in uncoupling transcription and translation of haploid male germ cells

**DOI:** 10.1101/2021.03.01.433315

**Authors:** Chong Tang, Mei Guo, Zhuoxing Shi, Zhuqing Wang, Chunhai Luo, Sheng Chen, Fengying Ruan, Zhichao Chen, Linfeng Yang, Xiongyi Wei, Chuanwen Wu, Bei Luo, Zhou Lv, Jin Huang, Dong Zhang, Cong Yu, Qiang Gao, Ying Zhang, Wei Yan, Fei Sun

**Author notes:** These authors contributed equally to this work: Chong Tang, Mei Guo, Zhuoxing Shi, Zhuqing Wang, and Chunhai Luo. Correspondence: Chong Tang,; Fei Sun,; Wei Yan,; Ying Zhang. Contact author: Wei Yan.

## Abstract

As one of the post-transcriptional regulatory mechanisms, transcription and translation’s uncoupling plays an essential role in development and adulthood physiology. However, it remains elusive how thousands of mRNAs get translationally silenced while stability is maintained for up to hours or even days before translation. In addition to oocytes and neurons, developing spermatids have significant uncoupling of transcription and translation for delayed translation. Therefore, spermiogenesis represents an excellent *in vivo* model for investigating the mechanism underlying uncoupled transcription and translation. Through full-length poly(A) deep sequencing, we discovered dynamic changes in poly(A) length through deadenylation and re-polyadenylation. Deadenylation appeared to be mediated by microRNAs (miRNAs), and transcripts with shorter poly(A) tails tend to be sequestered into ribonucleoproteins (RNPs) for translational repression and stabilization. In contrast, re-polyadenylation allows for translocation of the translationally repressed transcripts from RNPs to polysomes for translation. Overall, our data suggest that miRNA-dependent poly(A) length control represents a novel mechanism underlying uncoupled translation and transcription in haploid male germ cells.

## Introduction

Once synthesized, transcripts undergo extensive post-transcriptional modifications at both nucleus and cytoplasm [1]. In the nucleus, the premature mRNAs are processed into mature mRNAs by removing introns through splicing and adding 5’ caps and 3’ polyadenylated (poly(A)) tails. The poly(A) tail is critical for nuclear export, stability, and translation of mRNAs [2, 3]. In eukaryotic somatic cells, most cytoplasmic mRNAs’ poly(A) tails are shortened over time through deadenylation [4]. The shortening of poly(A) tail leads to reduced translational efficiency and increased degradation. Interestingly, the poly(A) tails of mRNAs can also be lengthened through cytoplasmic polyadenylation in specific cell types, including oocytes, early embryos, and neurons [5–7]. In mature oocytes, although mRNAs have shorter poly(A) tails (<20nt), they are stable and stored in ribonucleoprotein (RNP) particles without being translated [8, 9]. Soon after fertilization, these maternal transcripts are re-activated by cytoplasmic polyadenylation, which lengthens the poly(A) tails up to ∼80-150nt followed by efficient translation to produce proteins that are essential for the survival and growth of the embryos, from the fertilized egg to the stage that the zygotic genome is activated (2-cell embryo stage and 4-cell embryo stage in mice and humans, respectively) [9, 10]. The physiological significance of such a long delay in translation lies in that post-fertilization development before zygotic genome activation requires many proteins, which must be synthesized using pre-transcribed and stored maternal transcripts. In neurons, transcribed mRNAs tend to accumulate in the cell body, and these transcripts are sequestered into RNP granules, which travel a long distance and then start translation when reach the axon terminals [11–13]. Similarly, the translationally repressed mRNAs in neurons tend to have shorter poly(A) tails. Once they reach the synaptic junctions, these transcripts undergo cytoplasmic polyadenylation to lengthen their poly(A) tails, followed by an efficient translation [14, 15]. These findings suggest that ploy(A) length control represents an integral mechanism underlying uncoupled transcription and translation.

In addition to oocytes and nerve cells, haploid male germ cells, i.e., spermatids, also display uncoupled transcription and translation [16, 17]. As soon as round spermatids start to elongate, transcription is shut down due to the onset of nuclear condensation. However, from the onset of spermatid elongation (step 9 in mice) to the completion of spermatid differentiation into spermatozoa (step 16 in mice), there are numerous steps through which structurally sound spermatozoa are assembled [16, 18]. Since transcription ceases upon elongation (step 9), all proteins needed for the rest of the steps of sperm assembly (steps 9-16 in mice) have to be produced using transcripts pre-synthesized before the transcriptional shutdown, i.e., in round spermatids (steps 1-8) and even in late pachytene spermatocytes. For example, *Spata6* mRNAs start to be expressed in late pachytene spermatocytes, and its mRNA expression persists through the entire haploid phase. However, its protein is not detected until the sperm connecting piece (neck) starts to assemble in step 9 spermatids [19]. Therefore, spermiogenesis, the process through which round spermatids differentiate into elongated spermatids and eventually spermatozoa, represents an excellent *in vivo* model for studying uncoupling transcription and translation [20, 21].

Uncoupling of transcription and translation in spermiogenesis is achieved through physical sequestration of mRNAs subjected to translational delay into the RNP granules, which exist as the Nuage (also called intramitochondrial cement) in spermatocytes and the chromatoid body in round spermatids [18, 22]. When spermiogenesis progresses to elongation steps, these mRNAs are gradually released from RNPs and loaded onto polysomes to translate into the proteins required for sperm assembly [16, 18, 20, 21]. Recent works have shed light on the underlying molecular mechanisms. Briefly, our earlier work has revealed that RNP enrichment of mRNAs is a dynamic process, through which the overall length of 3’ UTRs becomes increasingly shortened compared to that of polysome-enriched mRNAs when late pachytene spermatocytes develop into the round and elongated spermatids [23]. The global 3’ UTR shortening is achieved through continuous shuffling of longer 3’ UTR mRNAs out of RNPs followed by UPF1-3-mediated, selective degradation [24] and by targeting shorter 3′ UTR mRNAs into RNPs [23]. In this way, the overall 3’ UTR length of the entire mRNA transcriptome in elongating spermatids becomes shorter and shorter [23]. We have also reported data showing that both miRNAs and m6A modification on pre-mRNAs are involved in the global shortening of transcripts and delay translation [25, 26]. Precisely, proper m6A levels control correct splicing and, consequently, the expected length distribution of transcripts [25]. Moreover, miRNAs target transcripts with longer 3′ UTRs through binding the distal binding sites to polysomes for translation followed by degradation, whereas transcripts with shorter 3′ UTRs only possess proximal miRNA binding sites, which, once bound by miRNAs, are targeted into RNPs for stability and translational repression [23].

Previous studies have shown that cytoplasmic poly(A) polymerases and poly(A) binding proteins are essential for spermiogenesis [27–30]. However, it remains unknown how poly(A) length is regulated during the global shortening of 3’ UTRs and the dynamic translocation of mRNAs between RNPs and polysomes during spermiogenesis, due to technical difficulties in determining the full-length sequences of the poly(A) tails. Although both TAIL-seq and PAL-seq have been developed as the next generation sequencing (NGS)-based methods for determining poly(A) tail sequences [31, 32], the short reads of NGS (<300nt) do not allow for accurate determination of full-length poly(A) sequences, thus compromising analyses on the relationship among 3’UTR length, poly(A) tail length, exon splicing patterns and translational status. To overcome this problem, we developed a sensitive method based on the third-generation PacBio sequencing, which we termed as <u>poly(A)</u>-<u>Pa</u>cBbio sequencing (PAPA-seq), similar to FLAM-seq[33]. PAPA-seq can accurately measure poly(A) length with reads covering the entire 3’ ends and the full-length transcripts. Using this method, we discovered that poly(A) length regulation pattern, through deadenylation and cytoplasmic re-polyadenylation, in differentiating spermatids. Moreover, we found that the miRNA is an ensential factor to regulate the poly(A) length and control the delay translation. For the first time, our data demonstrate the critical roles of poly(A) length and miRNA in uncoupling of transcription and translation, which is essential for normal spermiogenesis and male fertility.

## Results

### Dynamic changes in poly(A) length correlate with extended stability and delayed translation of mRNAs in developing haploid male germ cells

Given that the poly(A) tail is well-known to affect mRNA stability and translational efficiency [2, 3], we set out to measure the poly(A) length in spermatogenic cells using PAPA-seq (Figure 1A, 1B), a sensitive method similar to FLAM-seq [34]. To construct libraries for PAPA-seq, the poly(U) polymerase was used to add a poly(GI) tail to the native poly(A) tail, and the reverse transcription was then performed to generate cDNAs containing the full-length poly(A) tails followed by sequencing using the PacBio system (Figure 1B). Spike-in RNAs were sequenced to cross-validate the PAPA-seq data (**Supplemental Figure S1**). Using a modified STA-PUT method [23], pachytene spermatocytes, round and elongating spermatids were purified from adult mouse testes with purities of 90%, 90%, and 75%, respectively (Figure 1A, **Supplemental Figure S2**).

**Figure 1.**
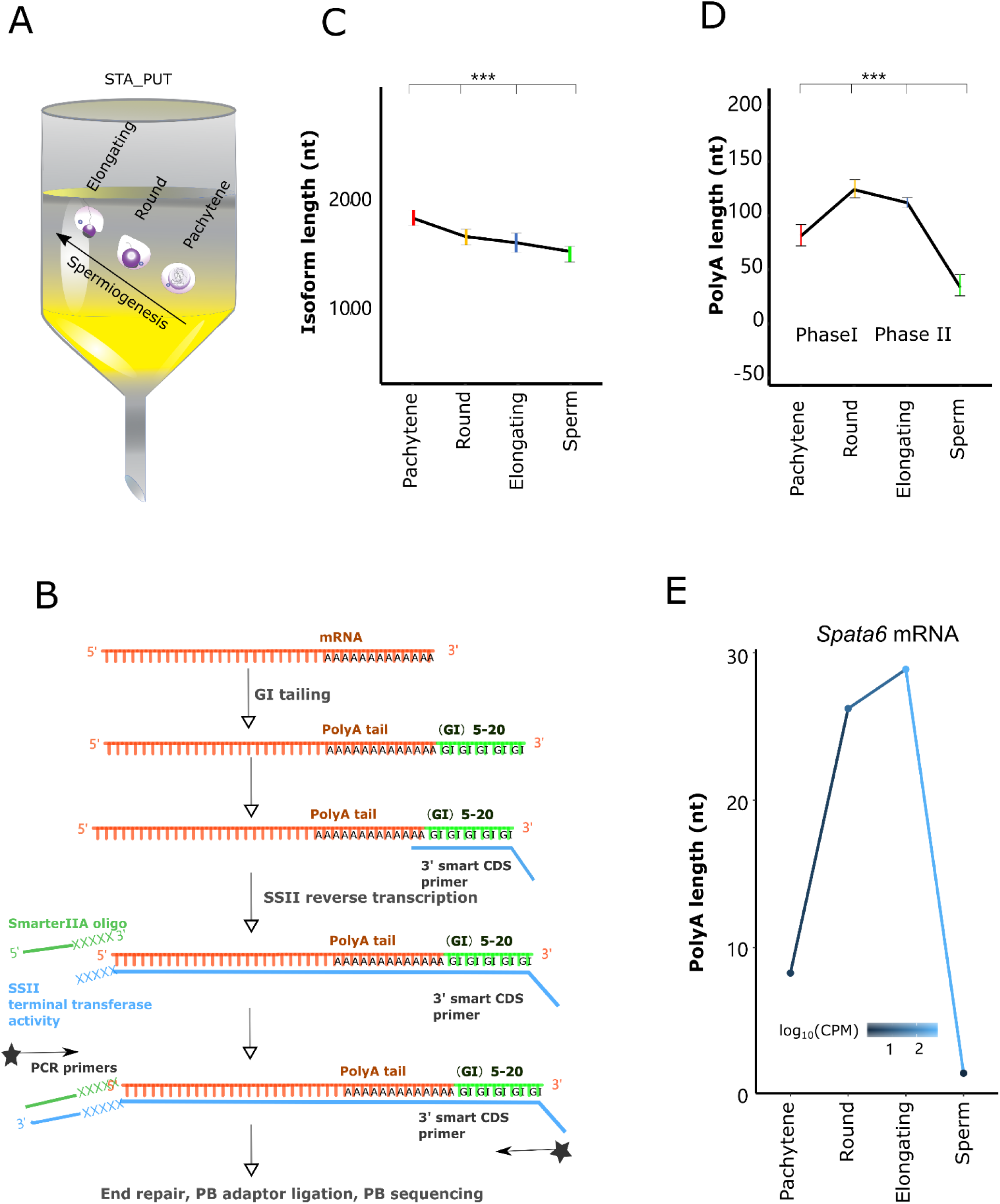
Dynamic poly(A) length control during spermiogenesis as revealed by full-length poly(A) deep sequencing. (A) A gravity sedimentation-based STA-PUT method used to purify pachytene spermatocytes, round spermatids and elongating spermatids in the present study. (B) Schematic illustration of PAPA-seq workflow. In brief, the poly(U) polymerase attaches poly(GI) tails to the very end of the poly(A) tail of RNAs. The modified poly(C) primer with adaptor sequence anneals to the poly(GI) tails. The reverse transcription, initiating from the start sites of poly(GI) tails, generates the cDNAs covering the full-length poly(A) tails. The other chemically modified adaptor is attached to the end of the cDNAs (i.e., corresponding to the 5’ ends of the RNA) by the template-switching activity of MMLV reverse transcriptase. PCR is then performed to amplify the cDNAs to a sufficient amount for PacBio library construction. (C) Average lengths of transcripts in pachytene spermatocytes, round and elongating spermatids, as well as spermatozoa. ***: p< 0.01, log t-test, number of transcripts >20,000. Data were based on samples from two independent preparations with 3-6 mice in each, plus two technical replicates. (D) Line plot showing the average poly(A) length in pachytene spermatocytes, round and elongating spermatids, as well as spermatozoa. ***: p< 0.01, log t-test, number of transcripts > 20,000. Data were based on samples from two independent preparations with 3-6 mice in each, plus two technical replicates. (E) Dynamic changes in the poly(A) length of *Spata6* transcripts during spermiogenesis. The x-axis stands for the male germ cell types, and the y-axis represents the average poly(A) length. CPM is indicated by lines with blue gradients. *Spata6* mRNA levels increase from pachytene spermatocytes to round spermatids and then peak in elongating spermatids while the poly(A) tails are lengthening during the same period.

Using PAPA-seq, we first examined the UTRs length during spermatogenesis. Consistent with our previous report by short reads sequencing [23], PAPA-seq data showed that the 3’ UTRs of mRNAs were progressively shortened when pachytene spermatocytes developed into the round and elongating spermatids (**Supplemental Figure S3A**). A similar trend was also observed in the 5’ UTRs, but to a lesser extent (**Supplemental Figure S3B**). Supporting our previous finding that m6A-dependent splicing activities increase with the progression of spermiogenesis [25], further analyses of the PAPA-seq data also revealed increased splicing events (alternative exon, exon skipping etc.) when pachytene spermatocytes developed into the round and then elongating spermatids (**Supplemental Figure S4A**). The global shortening of 3’ UTRs is believed to enhance translational efficiency because shorter 3’ UTRs provide fewer binding sites for regulatory factors, including RNA-binding proteins (RNPs) and small non-coding RNAs (sncRNAs) [23, 24, 35].

Although poly(A) length has long been known to influence mRNA stability and translational efficiency [2, 3], it remains unknown how the poly(A) length is regulated and whether the poly(A) length control is involved in uncoupling of transcription and translation during spermiogenesis. Despite the progressively shortened 3’ UTRs **(Supplemental Figure S3)** and decreased overall isoform transcript length (Figure 1C), we found that the poly(A) tail length was dynamically regulated in a biphasic fashion from pachytene spermatocytes to round and elongating spermatids (Figure 1D). The poly(A) tail length first increased from pachytene spermatocytes to round spermatids, which may contribute to the longer half-life of mRNAs that are pre-protected for delayed translation in late spermiogenesis (**Phase I**, Figure 1D **and Supplemental Figure S5**). In contrast, from round to elongating spermatids, the poly(A) length gradually decreased (**Phase II**, Figure 1D), suggesting that the shortening of the transcripts poly(A) tails coincides with the delayed translation progressing. Good agreement was found when comparing this result to the previous reports showing that shortening of the poly(A) tails correlates with translational activation in spermiogenesis [36, 37]. For example, transcript isoforms of *Spata6* mRNAs start to be expressed in late pachytene spermatocytes, and more isoforms continue to be expressed through the entire haploid phase. However, its protein is only expressed in the developing connecting piece in elongating (steps 9-12) and elongated spermatids (steps 13-16) [19]. We observed that the poly(A) length of *Spata6* drastically increased from pachytene spermatocytes to elongating spermatids, and this increase in poly(A) length occurred before the peak expression of SPATA6 protein in elongated spermatids/spermatozoa (Figure 1E). Taken together, our PAPA-seq analyses revealed that mRNAs in round spermatids owned the longest poly(A) tails compared to pachytene spermatocytes and elongating/elongated spermatids despite the global shortening trend in overall length of the isoform transcripts. The increased poly(A) length may function to enhance stability to support delayed translation during late spermiogenesis

### miRNAs mediate deadenylation of mRNAs enriched in RNPs

To further explore the effects of poly(A) length on mRNA translational repression and activation status, we fractionated cytosol of the three types of spermatogenic cells into RNP, mono-and poly-some fractions using sucrose gradient centrifugation followed by PAPA-seq. By measuring OD_254_, three fractions were observed: RNPs (Nuage/intramitochondrial cement in pachytene spermatocytes and chromatoid body in round spermatids), monoribosome, and polyribosome fractions (Figure 2A). The fractions’ purity was validated through the well-known markers distributed in RNP/polysome fractions (**Supplemental Figure S6**). Transcripts in the polysome fractions actively undergo translation, whereas those in the RNP factions are translationally suppressed [16, 20, 21, 38]. Interestingly, through PAPA-seq, we found that the average poly(A) length of RNP-enriched mRNAs was only 1/40 of those enriched in the polysome fractions (Figure 2B), suggesting that the deadenylation has a strong relationship with the localization to RNPs. If this is the case, then a question arises: how are the mRNAs selected for deadenylation followed by sequestration in RNPs? Previous studies have shown that small non-coding RNAs (sncRNAs) are enriched in RNPs [23] and that Argonaute proteins (e.g., AGO2, MIWI, and MIWI2) are abundant in the chromatoid body [39], suggesting the relationship of between sncRNAs and deadenylating the target mRNAs. Following these clues, we hypothesized that the enhanced deadenylation of the transcripts that subjected to translational suppression in the RNP phase during spermiogenesis might involve sncRNAs. To substantiate this hypothesis, we analyzed the miRNA-mRNA target relationship in RNPs and polysomes. We found that RNP-enriched miRNAs’ binding sites in the RNP-enriched transcripts with shorter poly(A) tails (∼5nt) were much more concentrated at the 3’ ends than those in polysome-enriched transcripts with longer poly(A) tails (∼200nt) in pachytene spermatocytes (Figure 2C), suggesting that miRNAs may target 3’ UTRs of the transcripts to initiate deadenylation, thus driving mRNA into RNPs. These results are in full agreement with previous studies about the miRNA driving mRNA into RNPs [23] and deadenylation [40–43].

**Figure 2.**
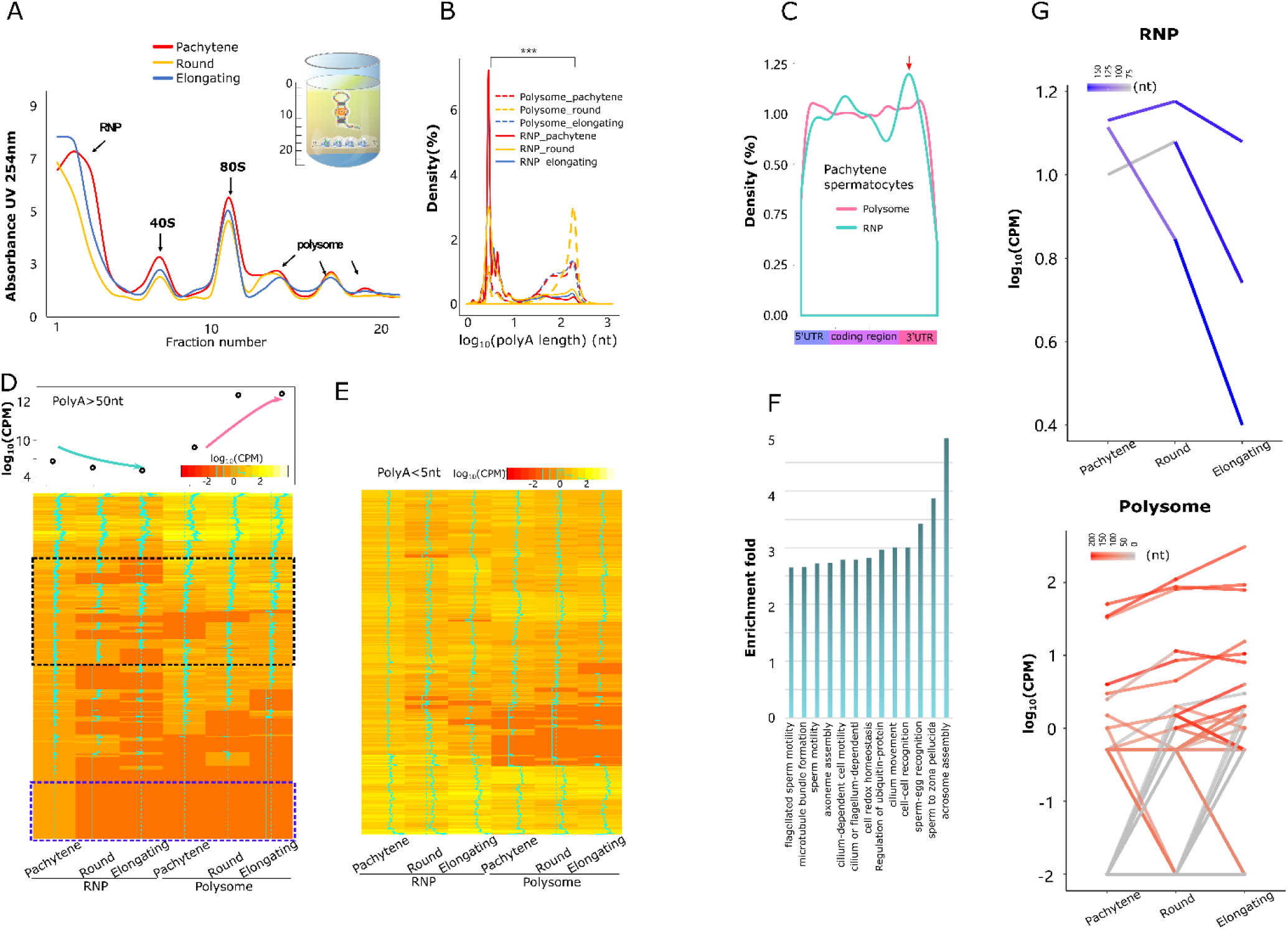
Poly(A) length distribution in the RNP granules and polysome fractions. (A) The sucrose gradient centrifugation separates the RNP granules and polysome fractions from purified pachytene spermatocytes, round and elongating spermatids. The diagram shows the RNA abundance (y-axis) in RNP, mono-and polysome fractions (x-axis). (B) Density plots showing poly(A) length distribution in the RNP granules and polysomes in pachytene spermatocytes, round and elongating spermatids. The RNAs with shorter poly(A) tails are enriched in the RNP granules with an average poly(A) length of 5 nt, whereas those with longer poly(A) tails are enriched in the polysome fractions with an average length of 200 nt. ***: p< 0.01, log t-test, number of transcript > 10,000. Data were based on samples from two independent preparations with 3-6 mice in each, plus two technical replicates. (C) Density plots showing distributions of the bioinformatically predicted miRNA targeting sites in transcripts enriched in RNP granules and polysome fractions in pachytene spermatocytes. The y-axis represents the density of the targeting sites, while the x-axis shows the full-length mRNAs. The RNP-enriched miRNAs preferentially target the 3’UTRs of mRNAs. (D) Heatmap showing that levels of mRNAs with the newly added longer poly(A) tails (>50bp) gradually decrease in RNP granules and increase in the corresponding polysome fractions in the three spermatogenic cell types. The top panel shows the average expression levels of these mRNAs in each fraction. (E) Heatmap showing that the levels of mRNAs with shorter poly(A) tails (<5nt) do not significantly change in RNP fractions. (F) GO term enrichment analyses of the mRNAs with longer poly(A) tails (>50nt) in the RNP granules of three male germ cell types. (G) Changes in expression levels of 25 *Spata6* isoforms in the RNP granules and polysomes. The x-axis indicates the three spermiogenic cell types, and the y-axis shows the levels/CPM of various isoforms. The specific color scheme corresponds to various poly(A) lengths. Levels of *Spata6* isoforms decrease with the increasing poly(A) length in RNP granules (darker blue indicates longer poly(A). In contrast, levels of *Spata6* isoforms increase with poly(A) lengthening (darker red indicates longer poly(A) in polysomes. Data were based on samples from two independent preparations with 3-6 mice in each, plus two technical replicates.

Generally, in somatic cells, the miRNA-based deadenylation triggers the targeted mRNA decaying [43]. In contrast, the chromatoid body (RNP) in spermiogenic cells functions to store transcripts [44] and the fate of the deadenylated transcripts stored in RNP is not limited to decaying. To address the fate of the deadenylated transcripts, we compared the poly(A) tail length distributions between RNP and polysome fractions in the three types of spermiogenic cells. Surprisingly, increased polyadenylation could be detected in both RNP and polysome fractions in round spermatids (**Supplemental Figure S7**). The presence of the newly polyadenylated tails in polysome fractions could mean extended stability during translation, whereas the newly polyadenylated transcripts in RNP phases may function to move transcripts out of RNPs and reach the polysomes, thus switching from translational suppression to active translation. To test this, we selected all of the transcripts with newly added poly(A) tails in RNPs (>50nt poly(A)) and examined their expression during spermiogenesis (Figure 2D **and Supplemental Figure S8**). Indeed, these transcripts’ levels decreased in RNP fractions but increased in polysome fractions from pachytene spermatocytes to round and elongating spermatids (Figure 2D**, significance in black square marked**). In contrast, the transcripts without a new poly(A) tail in RNPs (<5bp) stayed in the RNP fractions and remained translationally repressed (Figure 2E). A similar phenomenon was also observed in newly polyadenylated transcripts in round spermatid RNP fracrtions, in which the transcripts with longer poly(A) tail (>50nt) moved out of the RNP to polysome fractions, while those with shorter poly(A) tails (<5nt) remained in RNPs (**Supplemental Figure S8**). We also found that partial newly polyadenylated transcripts in RNPs quickly degraded in both RNP and polysome fractions (Figure 2D**, significance in blue squre marked**). These data suggested that the newly polyadenylated transcripts in the RNPs are either loaded onto the polysomes for active translation or subject to degradation after translation (Figure 2D **and Supplemental Figure S9**). GO term enrichment analyses revealed that those newly polyadenylated transcripts encode delayed translated proteins critical for sperm assembly and function, e.g., flagella development, acrosome assembly, motility, and sperm-egg recognition, etc.(Figure 2F). For example, *Spata6* is expressed as multiple isoforms during spermiogenesis and functions to form the sperm connecting piece/neck [19]. Once *Spata6* transcripts in the RNP phase gained the new long poly(A) tails (Figure 2G**, upper panel**), their levels dramatically decreased, whereas *Spata6* levels in polysome fractions were upregulated with a gradual increase in poly(A) tail length (Figure 2G**, lower panel**). By examining the terminal sequences of all of the RNP-enriched transcripts, we found that the transcripts without new poly(A) tails displayed ∼10x higher uridine contents than the transcripts with new poly(A) tails (**Supplemental Figure S10D**), implying the function of the uridine-rich motif in polyadenylation. Taken together, our data suggested that sncRNAs, especially miRNAs, likely mediate deadenylation of transcripts to be sequestered in RNPs, and only the mRNAs with shorter poly(A) tails can be sequestered in RNPs. Moreover, gaining longer poly(A) tails is a prerequisite for translocation from RNPs to polysomes for translation.

### miRNA ablation causes a failure in both poly(A) shortening and RNP phase separation in developing spermatids

To further justify the aforementioned proposal, we first employed the *Drosha* cKO mouse model. *Drosha* encodes a nuclear RNase III enzyme essential for pre-miRNA cleavage [45], and inactivation of *Drosha* exclusively in the spermatogenic cell linage through a conditional knockout (cKO) approach can abolish miRNA production in all developing male germ cells [46]. Although *Drosha* cKO testes contain fewer spermatogenic cells due to germ cell depletion, there are still some pachytene spermatocytes, and round spermatids remained in the seminiferous epithelium [46], which we purified and pooled for PAPA-seq analyses.

To further validate our notion that miRNAs bind to the 3’ UTRs of their target mRNAs to deadenylate and sequester mRNAs into RNPs, we analyzed *Drosha*-null spermatogenic cells and examined the effects of miRNA deficiency on the poly(A) length of their target mRNAs. RNA contents in the RNP fractions of the *Drosha*-null pachytene spermatocytes and round spermatids were significantly lower than those in the wild-type counterparts (Figure 2A **and** 3A), suggesting that these transcripts might fail to accumulate in the RNP granules in the *Drosha* cKO male germ cells. Moreover, the ratio of gene numbers in RNP vs. polysome fractions in *Drosha*-null spermatogenic cells decreased by >2 folds when compared to that in wild-type spermatogenic cells (Figure 3B), further suggesting that in *Drosha-*null male germ cells, the transcripts failed to be compartmentalized to RNP granules in the absence of miRNAs. The fact that the mRNA levels in RNP fractions of *Drosha* cKO cells were drastically reduced compared to those in RNP fractions of wild-type cells (Figure 3C) also seems to support this possibility because the transcripts that failed to be localized to RNPs undergo massive degradation, leading to much-reduced RNA levels. The results agree with the phenotype that we observed in the *Drosha* cKO mice, showing that ablation of miRNAs disturbed the delay translation in spermiogenesis [47]. Further analysis of the poly(A) in different cell components showed that the average poly(A) length of RNP-enriched transcripts in *Drosha-*null male germ cells was significantly longer than those in wild-type cells (Figure 3D **and Supplemental Figure S11**), supporting the notion that miRNAs function to trim poly(A) tails. The increased average poly(A) length in RNP-enriched transcripts in the *Drosha*-null germ cells most likely resulted from relative enrichment of longer poly(A) transcripts due to the failed compartmentalization of the transcripts with trimmed poly(A) tails into the RNP granules in the absence of miRNAs. To further support this notion, we examined the dynamic poly(A) change between RNPs and polysomes in the wild-type and the *Drosha*-null round spermatids (Figure 3E). We found that these *Drosha*-null transcripts failed to be sequestered into RNPs, and appeared to be stuck in the polysome fractions with very long poly(A) tails (Figure 3E). It is also noteworthy that the partial RNP-enriched transcripts in the wild-type round spermatids possessed much shorter poly(A) tails, and the same set of transcripts were mostly stuck in *Drosha*-null RNPs with much longer poly(A) tails (Figure 3E**, significance in black square marked**. The miRNA influence on the deadenylation of transcripts and translocation of the transcripts, is pertinent. For example, the multiple isoforms of *Spata6* were tailed with longer poly(A) in polysomes and shorter poly(A) in RNP granules in wild-type round spermatids, respectively (Figure 3F). In sharp contrast, most of these transcripts were degraded in *Drosha* cKO cells, likely due to mRNA degradation in the absence of miRNAs (Figure 3F). As expected, a few transcripts with much longer poly(A) tails remained in the polysome fractions in *Drosha* cKO cells (Figure 3F). Taken together, our data suggest that miRNAs stabilize and facilitate compartmentalization of their target transcripts to the RNP granules, and miRNA depletion causes their targets to degrade without being compartmentalized into RNPs. In *Drosha*-null spermatogenic cells, miRNA deficiency failed to shorten the poly(A) tails of their target RNAs timely, leading to a degradation or a retention in polysome rather than compartmentalization into the RNPs.

**Figure 3.**
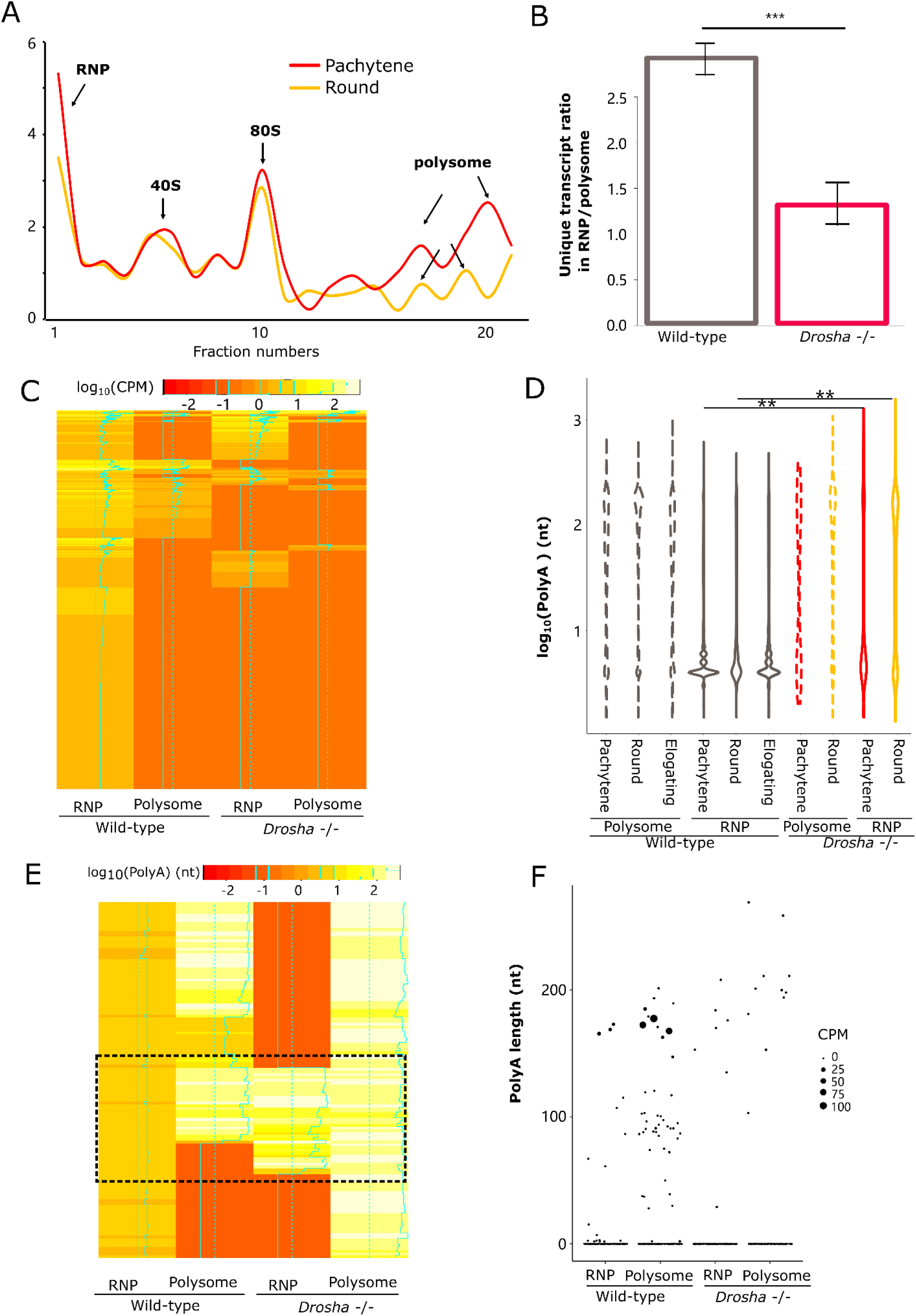
Effects of miRNA deficiency on the poly(A) length distribution in the RNP granules and polysome fractions. (A) Distribution of RNA contents in the RNP and polysome fractions in purified *Drosha*-null pachytene spermatocytes and round spermatids. (B) Bar graphs showing the ratio of RNP-enriched vs. polysomes-enriched transcripts in wild-type and *Drosha*-null pachytene spermatocytes and round spermatids. ***: p< 0.01, Wilcoxon rank test. Wild-type samples were from two independent preparations with 3-6 mice in each, plus two technical replicates. *Drosha* KO samples were from single preparation with 6-10 KO testes plus two technical replicates. (C) Heatmap showing the expression level of wild-type RNPs enriched transcripts among fractions. RNP enriched transcripts in wild type round spermatids are generally down-regulated in both RNP and polysome fractions in *Drosha*-null round spermatids. (D) Violin plots showing the poly(A) length distribution in the RNP and polysome fractions of wild-type and *Drosha*-null spermatogenic cells. **: p< 0.05, Student t-test, number of transcripts > 10,000. Wild-type samples were from two independent preparations with 3-6 mice in each, plus two technical replicates. *Drosha* KO samples were from single preparation with 6-10 KO testes plus two technical replicates. (E) Heatmap showing poly(A) length of transcripts enriched in RNP in wild-type round spermatids are largely absent in the RNPs of *Drosha*-null round spermatids, and these appear to be stuck in the polysome fractions with elongated long poly(A) tails. Partial transcripts also could bind to RNPs with long poly(A) tails (Black square). (F) Dot plot showing the distribution of the abundance of *Spata6* isoforms with different poly(A) length in both RNP and polysome fractions of wild type and *Drosha-*null round spermatids. Wild-type samples were from two independent preparations with 3-6 mice in each, plus two technical replicates. *Drosha* KO samples were from single preparation with 6-10 KO testes plus two technical replicates.

### X-linked *miR-506* family miRNAs sequester *Fmr1* mRNAs into RNPs after deadenylation

In our proposed model, mRNAs need to shorten their poly(A) tails through miRNAs-mediated deadenylation to be sequestered into RNP granules for delayed translation. However, the *Drosha* cKO model is more likely to experience analysis problems in systematical mRNA destabilization caused by massive miRNA changes. Therefore, we chose to utilize the *miR-506* family knockout mouse line that we generated [48] to investigate the effects of ablation of 18 miRNAs on their target mRNA, *Fmr1*, emphasizing the poly(A) length and translocation between RNPs and polysomes. The relationship between poly(A) and miRNAs may be reflected more precisely by utilizing the single miRNA family knockout model.

The *miR-506* family contains 21 miRNAs transcribed from five large miRNA clusters encompassing a ∼ 62kb region and a ∼22kb region near *Slitrk2* and *Fmr1*, respectively, on the X chromosome, most of which are preferentially expressed in the testis [48]. The KO line used in this study lack 18 abundantly expressed miRNAs out of the 21 miRNAs that belong to the *miR-506* family [48]. The *miR-506* family targeting *Fmr1*, has been validated using Western blot *in vivo* [48] and Western blot as well as luciferase assays *in vitro* [49] (Figure 4A). We first analyzed *Fmr1* mRNA levels using a semi-quantitative PCR and found no changes in *Fmr1* levels in KO testes (Figure 4B), indicating that *miR-506* did not degrade its target *Fmr1*. We then determined levels of FMRP, the protein encoded by *Fmr1*, using Western blots (Figure 4C). To our surprise, FMRP levels were reduced by ∼50% in the *miR-506* family KO testes compared to the wide-type controls (Figure 4C). To unveil the differential patterns between mRNA and protein, we further examined the poly(A) length in the *miR-506* family KO and WT testes using a poly(A) length PCR assay kit, as described previously [50]. Interestingly, the average poly(A) length of the *Fmr1* mRNAs appeared to be doubled in the *miR-506* KO testes (∼420nt in WT and ∼800nt in KO testes) (Figure 4D), suggesting that the poly(A) length affected protein translation effeciency. The poly(A) pattern of the *miR-506* family KO mice is consistent with that of the *Drosha* cKO mice, supporting that the functional roles of miRNAs in the testis is to trim poly(A) tails rather than degrading their targets. Based on our proposed model, the *Fmr1* mRNA would not be able to be deadenylated and sequestered into RNP granules in the absence of *miR-506* family miRNAs. Indeed, our data revealed that *Fmr1* mRNAs levels decreased by ∼10% in RNP granules but increased in the polysome fractions (Figure 4E **and Supplemental Figure S12**). Moreover, the poly(A) tails of *Fmr1* mRNAs in RNPs are much longer in the *miR-506* family KO testes than those in the wild-type testis (Figure 4F), and no significant difference were observed in the polysome fractions (Figure 4E). These findings are generally consistent with those found in the *Drosha* cKO testes, where some mRNAs with longer poly(A) tails still could bind to RNP granules despite massive miRNA depletion (Figure 3E **and** Figure 3F). Overall, these results strongly support our hypothesis that miRNAs shorten the poly(A) length of their target mRNAs by deadenylation to sequester their target mRNAs into RNP granules for delayed translation.

**Figure 4.**
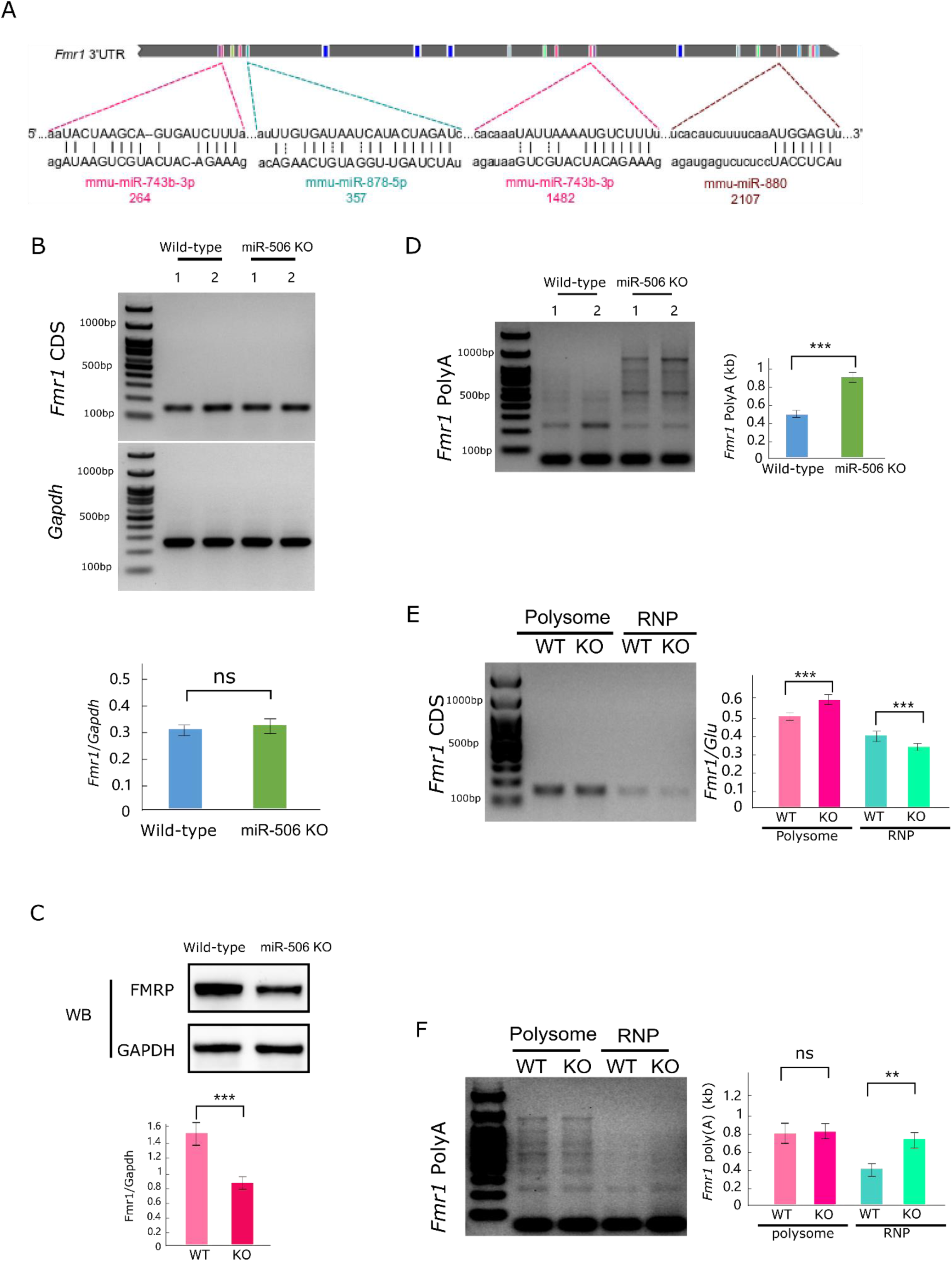
*miR-506* family miRNAs regulate the poly(A) length of their target gene *Fmr1* in the testis. (A) Schematic illustration showing the targeting sites in *Fmr1* 3’ UTR by 4 of the *miR-506* family, as previously reported [49]. (B) Semi-quantitative RT-PCR analyses on levels of *Fmr1* mRNA (CDS for coding sequences) in total testes of the *miR-506* family KO and wild-type male mice. *Gaphd* was used as the loading control. Histograms display quantitative analyses of the data (mean ± SEM) from biological triplicates (n=3). (C) Western blot analyses of *Fmr1* protein (FMRP) levels in the *miR-506* family KO and wild-type testes. Data are presented as mean ± SEM, using biological triplicates (n=3). (D) Distribution of the poly(A) length of *Fmr1* mRNAs in total testes of the *miR-506* family KO and wild-type control male mice. Left panels show representative gel images, whereas right panels display quantitative analyses of the data (mean ± SEM) from biological triplicates (n=3). (E) Semi-quantitative RT-PCR analyses on levels of *Fmr1* mRNA (CDS for coding sequences) in RNP and polysome fractions in the *miR-506* family KO and wild-type testes. *Clu*, known not to display delayed translation was used as the loading control. The left panels show representative gel images. Histograms (right panels) display quantitative analyses of the data (mean ± SEM) from biological triplicates (n=3). (F) Distribution of the poly(A) length of *Fmr1* mRNAs in testicular RNP and polysome fractions of the *miR-506* family KO and wild-type control male mice. Left panels show representative gel images, whereas right panels display quantitative analyses of the data (mean ± SEM) from biological triplicates (n=3).

## Discussion

Uncoupling of transcription and translation is prominent during spermiogenesis (round spermatid differentiation into spermatozoa) [16, 20, 21], oogenesis (maternal transcript production) [51, 52], preimplantation embryonic development (protein production before zygotic genome activation) [9, 10, 53], and neuronal cell functions (mRNA synthesis in the cell body and translation in the axon) [11–13]. Several potential mechanisms have been identified to achieve uncoupled transcription and translation, including physical sequestration of mRNAs and proteins in RNP granules [20, 21, 23], 3’ UTR length control through alternative polyadenylation [54, 55], and 3’ UTR length-dependent, selective decay of transcripts by UPF proteins [24, 56]. The poly(A) length has long been known to regulate transcript stability and translational efficiency [2, 3]. However, investigations into the role of poly(A) length control have just started to emerge [2, 8, 34, 57]. This is primarily due to a lack of sensitive methodologies that allow for accurate determination of the full-length poly(A) tail sequences. The third-generation deep sequencing technologies, e.g., the PacBio and Nanopore sequencing, allowed us to develop a sensitive method, which we termed PAPA-seq, to determine the full-length sequences of not only poly(A) tails but also the rest of the entire transcripts. Using PAPA-seq, we discovered that the average length of the ployA tails is ∼100 nt. However, it can be as long as 1,000nt in some transcripts in the three spermatogenic cell types, i.e. the pachytene spermatocytes, and the round and elongating spermatids. The poly(A) tails are much longer in the round and elongating spermatids than those in the pachytene spermatocytes. This pattern aligns well with the highest number of transcripts subjected to delayed translation in the round and elongating spermatids, compared to the pachytene spermatocytes. Since longer poly(A) tails tend to have enhanced stability and translational efficiency, the peak of poly(A) lengthening in the round and elongating spermatids may reflect the peak of the delayed translation of those pre-synthesized, RNP-enriched transcripts.

Previous reports have shown that PABPs interact with the deadenylation complex (CCR4-NOT-Tob and PAN2-PAN3) to cause increased mRNA decay and repressed translation by shortening their poly(A) tails [58, 59]. However, this mechanism targets all transcripts, causing massive degradation, whereas in developing spermatids with uncoupled transcription and translation, deadenylation occurs in the transcripts to be sequestered into the RNP granules. The selective deadenylation must be mediated by a factor with sequence specificity, e.g., miRNAs. Indeed, our data strongly support such a role of miRNAs. Therefore, miRNAs appear to play a important role by regulating the poly(A) length through binding their target mRNAs.

In the absence of miRNAs, the mRNAs with pure poly(A) tails cannot be deadenylated and thus, failed to be compartmentalized into the RNP granules, however, a few transcripts still bound with RNPs, which may due to other targeting sncRNAs and/or poly(A) patterns. In spermatogenic cells, the miRNA only counts for 10% of total sncRNAs [23]. Other sncRNAs, for example, tsRNA and piRNA, also might sequester mRNAs into RNPs [60, 61]. It is also noteworthy that the molar ratio between sncRNAs and large RNAs in RNPs is over 100:1, implying that the excessive amount of sncRNAs may target other regions of mRNAs to translocate mRNAs [23]. On the other hand, we also examine the poly(A) tail nucleotide distribution patterns. The cytosine-enriched poly(A) tails still can manage to be phase-separated into the RNP granules because the cytosine-enriched poly(A) tails have much reduced PABP binding affinity when compared to the pure poly(A) tails [62] and thus, are more capable of changing their internal structures to increase hydrophobicity, driving RNP phase separation independent of miRNA-mediated deadenylation. Overall, the miRNA binding is critical but not the only factor to sequester the transcripts into RNPs. Other factors, such as RNA binding proteins, non-A nucleotide distribution in poly(A) tails, and other sncRNAs, could also affect the mRNAs storage in RNPs.

Our earlier work has demonstrated that *Drosha* cKO testes display severe spermatogenic cell depletion with a few remaining pachytene spermatocytes and spermatids [47]. Therefore, the results observed in *Drosha* KO cells may represent a secondary effect, that could compromise our conclusion on the regulation of poly(A) length by miRNAs (Figure 3). To further validate our proposed model on the miRNA-mediated deadenylation and the subsequent subcellular compartmentalization (i.e. RNPs *vs.* polysomes), we employed another KO mouse line, which lacks five large miRNA clusters of the *miR-506* family (Figure 4). These KO males are subfertile, and their testes display normal spermatogenesis and histologically indistinguishable from the wild-type control testes [48]. The lack of germ cell depletion makes this KO mouse line advantageous due to minimal secondary effects. Interestingly, we found inconsistent levels between *Fmr1* mRNA and protein (Figure 4A **and** 4B). Specifically, *Fmr1* protein FMRP levels decrease by half, whereas its mRNA levels remain unchanged despite their poly(A) tails double their length in the absence of 18 of *Fmr1*-targeting miRNAs in the KO testes *in vivo* (Figure 4). This finding is surprising because it is against the common belief that miRNA degrade the target mRNAs in somatic cells [63]. However, this result does support our proposed model in spermiogenesis. Specifically, in the absence of targeting miRNAs, the mRNAs fail to be deadenylated. Although *Fmr1* mRNAs managed to get into RNPs, their translation in elongating and elongated spermatids does not occur normally, leading to a decreased protein level, because of the dysregulated poly(A) tails. This is in sharp contrast to the wild-type situation that *Fmr1* with shorter poly(A) tails are stored in RNPs. Taken together, our data support the notion that miRNAs triggers the deadenylation of their targeted mRNAs, and deadenylation of mRNAs effectively controls the delayed protein translation in spermiogenesis.

The function of the chromatoid body (RNPs) is still a source of debate, but the generally accepted theory is that the chromatoid body (RNPs) determine the transcript fate for degradation or storage for delayed translation [64–72]. Disturbing the chromatid body components caused severe defects during the spermiogenesis [73]. However, a definitive link between poly(A) tails and the dynamic changes between RNP and polysome has been challenging to prove. In our research, there appears to be a clear poly(A) length association between the RNP and polysome fractions (Figure 2). One of our most intriguing findings is that the RNP-stored transcripts were re-adenylated and moved to polysome for delayed translation in later developmental stages (Figure 2). This finding is a reasonable implication of the delayed translation model in spermiogenesis. However, the study is limited by the difficulty in distinguishing the early stored transcripted RNAs from the nascent RNAs in the re-polyadenylated RNAs population, considering the active translation in round spermatids. To support our notion, we refocused our analysis on the newly re-polyadenylated RNAs in the round spermatids (Supplemental Figure S8). Among these transcripts, the mRNA translocating phenomena are also observed in the elongating spermatids, in which the transcription is mostly ceased and the transcripts can only move from the RNPs (Supplemental Figure S8). Moreover, we also analyzed the ATAC-seq data to observe the transcription status [74]. Most of the re-polyadenylated mRNAs’ transcription regions are already condensed in the round and elongating spermatids, supporting that most of the RNAs are translocated from RNPs other than the nascently transcribed mRNA from nucleus. On the other hand, we also found that half of the transcripts bond in RNPs are degraded quickly. There is the dual mechanisms to select the transcripts to degrade or to store for delayed translation. Further researches need to locate the enzyme responsible for the re-polyadenylation and selective mechanism of the polyadenylation.

## Conclusion

In summary, we here reported, for the first time, the dynamic changes in poly(A) length and a critical role of miRNA mediate poly(A) control in the regulation of uncoupled transcription and translation in the male germ cells undergoing spermiogenesis.

## Methods

### Animals

All wild-type and KO mice used in this study were on the C57BL/6J background and housed under specific pathogen-free conditions in a temperature- and humidity-controlled animal facility at the Nantong University and University of Nevada, Reno, respectively. The animal protocols was approved by the Animal Care and Use Committee of Nantong University or University of Nevada, Reno. Male germ cell-specific *Drosha* conditional KO mice (*Drosha^loxp/loxp^* mice were bred with *Stra8-iCre* mice) and global KO of X linked *miR-506* family were generated and genotyped at the University of Nevada, Reno as previously described [23, 46, 48, 75].

### Purification of spermatogenic cells

Pachytene spermatocytes, round, and elongating/elongated spermatids were purified from adult mouse testes using the STA-PUT method [23]. The BSA gradients (0.5-4%) were prepared in the EKRB buffer (Cat#K-4002, Sigma), supplemented with sodium bicarbonate (1.26g per 1L), L-glutamine (0.29228g per 1L), Penicillin and Streptomycin mix (Thermo-Fisher, 10,000U per 1L), MEM non-essential amino acids (Thermo-Fisher, 1ml 100X per 1L), MEM amino acids (20ml 50X per 1L) and cycloheximide (100ng/ml), pH7.2-7.3). Eight testes were pooled each time for cell purification. After being collected and decapsulated, testes were placed into 10ml of the EKRB buffer containing 5mg collagenase (Sigma) for 12-min digestion at 32°C to disperse the testicular cells. Once dispersed, the testicular cells were washed three times using the EKRB buffer followed by trypsin digestion by incubation in 10ml EKR buffer containing trypsin (Sigma, 0.25mg/ml) and DNase I (Sigma, 20 μg/ml) at 37°C for 12min with occasional pipetting to facilitate cell dispersion. Thoroughly dispersed testicular cells were washed three times, followed by centrifugation and re-suspension in 10 ml of 0.5% BSA. The cell suspension was passed through 50 μm filters, and the filtrate was saved for loading onto the STA-PUT apparatus for sedimentation. After 3 h sedimentation at 4 °C, fractions were collected from the bottom of the sedimentation chamber. A total of 30 fractions of 15 ml each were collected. After centrifugation, the supernatants were removed, and the cells in each fraction were re-suspended and the cell purity was determined by microscopy examination based on cell morphology, as described previously [76]. Fractions containing the same cell types were pooled followed by centrifugation to collect purified pachytene spermatocytes, round spermatids, and elongating/elongated spermatids.

### RNP and polysome fractionation

We fractionated the purified spermatogenic cells into RNP, monoribosome, and polyribosome fractions using a continuous sucrose gradient ultracentrifugation method, as described [23]. In brief, a continuous sucrose gradient (15%-50%) was prepared by carefully overlaying 15% sucrose onto 50% sucrose followed by diffusing for 3 hours at 4°C. The 15% and 50% sucrose solutions were prepared in a lysis buffer (containing 150mM potassium acetate, 5mM magnesium acetate, 2mM DTT, protease inhibitor cocktail (Sigma, 1X), RNase inhibitor cocktail (Sigma, 1X), cycloheximide (100ng/ml), and 50mM HEPES, pH 7.5). Freshly purified pachytene spermatocytes, round spermatids, and elongating/elongated spermatids were homogenized in the lysis buffer freshly supplemented with 0.5% Triton X-100 and 0.25 M sucrose. The homogenates were centrifuged at ∼500g for 15min at 4°C to remove tissue debris, unbroken cells, and nuclei. The supernatant was loaded onto the continuous 15-50% sucrose gradient followed by centrifugation at 150,000g (35,000rpm) for 3h at 4°C. A tiny hole was punched gently at the bottom of the tubes for fraction collection. Twenty-four 500-ul fractions were collected followed by UV spectrometer measurement for OD254.

### PAPA-Seq

Total RNA from all samples (cell sample RIN>8, polysome sample RIN>8, RNP not applicable, sperm sample RIN>3) was centrifuged to discard inhibitor pellet. mRNA was purified using Dynabeads® mRNA Purification Kit (life technology) according to the manufacturer’s instruction. mRNA was checked by 2100 Bioanalyzer RNA picochip to ensure integrity. Qualified mRNA was firstly denatured at 65℃ and chill on ice, and then added with 1× GTP-ITP mix(0.5 mM each) and 1× NEB buffer 2, 2U poly(U) polymerase, and 40U RNase inhibitor, followed by incubation at 37℃ for 1hour. The mRNAs tailed with GTP and ITPs was then purified by 1.8X volumes RNA cleanup Ampure beads (Catalog NO. A63987, Beckmann Coulter) and eluted in 10.5μl H2O, followed by reverse transcription. 2μl 3’CDS primer (10μM) (5’-AAGCAGTGGTATCAACGCAGAGTACNNNNNNCCCCCCCCCCCCTTT-3’) was added into the purified GI-tailed mRNA, and incubated at 72℃ for 3min, followed by cooling down to 42 ℃ at the 0.1 ℃/s speed, then a master mix containing iso-template switch oligo (5’-AAGCAGTGGTATCAACGCAGAGTACATrGrG+G-3’, where +G indicates locked nucleotide), SSII superscript transcriptase, 1× first strand buffer, DTT, dNTP, and RNase inhibitor was added. The reaction was incubated at 42℃ for 90 minutes, and stopped by incubating at 70℃ for 10 minutes. The full-length mRNA library reaction was set up with the following reagents: 1 × KAPA HiFi mater mix, PCR primers (5’-AAGCAGTGGTATCAACGCAGAGT-3’), and 10 μl of synthesized cDNA. The PCR conditions were as follows: initial denaturation at 95°C for 2 min, followed by 15-18 cycles of amplification (denaturation at 98°C for 20 s, annealing at 65°C for 15 s, and elongation 72°C for 4 min) with a final elongation at 72°C for 7 min. The amplified library was then purified with 1 × and 0.4 × Ampure XP DNA Beads separately and resuspended in 42 μl H2O. One microliter of the equal mass mixed library was diluted 5 times and one microliter was checked on High sensitivity DNA chip.

### PacBio sequencing

The purified PCR libraries were submitted to the Genomics core facility of MDC for PacBio sequencing. Sequencing libraries were prepared using the PacBio Amplicon Template Preparation and Sequencing Protocol (PN 100-081-600) and the SMRTbell Template Prep Kit 1.0-SPv3 according to the manufacturer’s guidelines. Sequencing on the Sequel was performed in Diffusion mode using the Sequel Binding and Internal Ctrl Kit 2.0. Every library was sequenced on one SMRT Cells 1 M v.2.1 with a 1 × 1200 min movie. Circular Consensus Sequence (CCS) reads were generated within the SMRT Link browser 5.0 (minimum full pass of three and minimum predicted accuracy of 90).

### Bioinformatic analyses of PAPA-Seq data

#### Full-Length Isoforms detection

First, we used NCBI BLAST (the version is 2.2.28+ with parameters “-outfmt 7 -word_size 5”) to map 5’ and 3’ primers to CCS reads, then used in house Perl script to parse standard pair of 5’ and 3’ primers CCS as the full-length isoform. Next, we trimed the primer sequence and reported the UMI in each full-length isoform. Finally, each isoform was oriented from 5’ to 3’ end.

#### Poly(A) tails detection

We developed a special modified sliding window algorithmic approaches to accurately and error-tolerantly detect Poly(A) tails. For example, we have a following poly(A) tail: TCGAAATCAAGAAAAACAAAAAA, we listed all the windows without overlap (from 3’ to 5’: AAAAAA, AAAAAC, AAG, AAATC, TCG), and obtained the percentage of A in each window (100%, 83.33%, 66.66%, 60%, 0%), which were defined as the parameter w1. Then using sliding window started from the 3’ end, we can get the percentage of A in total window, which were defined as the parameter w2. Based on the empirically optimized and benchmarked against a set of manually annotated poly(A) tail estimated from UHRR datasets, the w1 and w2 parameters were set to w1>=50% and w2>=70%. As for the example above, we listed all the sliding region here (AAAAAA w1=100% w2=100%, AAAAACAAAAAA w1=83.33% w2=91.66%, AAGAAAAACAAAAAA w1=66.66% w2=80%, AAATCAAGAAAAACAAAAAA w1=60% w2=80%, TCGAAATCAAGAAAAACAAAAAA w1=0% w2=69.56%), so we can define the poly(A) is AAATCAAGAAAAACAAAAAA.

#### Quantification and gene assignment

After Poly(A) tails detection and trimming, the remaining fraction of each isoform was mapped to mm10 genome using GMAP (version is 2018-05-30) with parameters ‘-f samse -n 0 --min-intronlength 9 --max-intronlength-middle 500000 --max-intron length-ends 10000 --trim-end-exons 12’. And then using cDNA_Cupcake (https://github.com/Magdoll/cDNA_Cupcake) python script collapse_isoforms_by_sam.py to collapse all samples isoforms, based on collapsed output, we are using in house Perl script to get the isoform expression quantity in each sample. After collapse, nonredundant isoforms were detected using cuffcompare (version v2.0.2) assigned to Ensemble mm10 annotation gene models.

#### Isoforms coding frame prediction and UTR detection

CDS coding frame and UTR region were predicted using TransDecoder [77] form the nonredundant isoforms, the predicted CDS were further confirmed using NR and Pfam database.

#### In silico miRNA target prediction

Computational prediction of miRNA targets is a critical initial step in identifying miRNA: mRNA target interactions for experimental validation. In order to find possible targets, multiple software was used. The intersection targets with appropriate filter conditions such as MFE scores were taking for the further analysis. We used miRanda [78] (with parameters’ -en -20 - strict’) and TargetScan [79] (with default parameter) to get the target genes of miRNA, extracted intersection or union of the target genes as final prediction result.

#### Statistical analyses

Both student’s t-test and Wilcoxon rank sum test (a non-parametric or distribution-free test) were used for statistical analyses. The majority of the data followed a lognormal distribution. The student’s t-test was also performed on the logarithm data.

## Declarations

## Acknowledgements

This work was supported by grants from the National Key Research and Development Program of China (No. 2018YFC1003500 to F.S), the National Natural Science Foundation of China (Grant No.81430027 and 81671510 to F.S.), the Science, Technology and Innovation Commission of Shenzhen Municipality (No: JSGG20170824152728492). The work on *Drosha* and X linked miR-506 family KO mice were supported by grants from the NIH (HD071736 and HD085506 to WY) and the Templeton Foundation (PID: 61174 to WY). The funders had no role in the design of the study, data collection, analysis, and interpretation, or in writing the manuscript.

## Funding

This work was supported by grants from the National Key Research and Development Program of China (No. 2018YFC1003500 to F.S), the National Natural Science Foundation of China (Grant No.81430027 and 81671510 to F.S.), the Science, Technology, and Innovation Commission of Shenzhen Municipality (No: JSGG20170824152728492), National Natural Science Foundation of China Grants (Grant No. 81801523 to Ying Zhang). The work on Drosha and X linked miR-506 family KO mice was supported by grants from the NIH (HD071736 and HD085506 to WY) and the Templeton Foundation (PID: 61174 to WY). The funders had no role in the design of the study, data collection, analysis, and interpretation, or in writing the manuscript.

## Availability of Data and Materials

The original data of both large and small RNA-seq data have been deposited into the CNGB database and can be accessed using the accession numbers CNS0189582. The code has been uploaded to GitHub (https://github.com/shizhuoxing/BGI-Full-Length-RNA-Analysis-Pipeline).

## Author contributions

C.T., W. Y. F. S. and Y.Z supervise the entire study. Y.Z. and C.L. performed cell purification, fractionation, M.G. performed RNA isolation, library construction and sequencing. Z.W. and S.C. performed all the experiments involving the X linked *miR-506* family KO mice. C.T. and Z.S. conducted all the bioinformatics analyses. W.Y. provided purified germ cells from the *Drosha* cKO and the X linked *miR-506* family knockout mice. All participated in data analyses. C.T. and W.Y. wrote the manuscript. Z.W. edited the manuscript. All authors read and approved the final manuscript.

## Competing interest

The authors declare no competing interests.

Ethical approval and consent to participate

## Notes

### Competing Interest Statement

The authors have declared no competing interest.

